# Karyon: a computational framework for the diagnosis of hybrids, aneuploids, and other non-standard architectures in genome assemblies

**DOI:** 10.1101/2021.05.23.445324

**Authors:** Miguel A. Naranjo-Ortiz, Manu Molina, Verónica Mixão, Toni Gabaldón

**Affiliations:** Centre for Genomic Regulation (CRG), The Barcelona Institute of Science and Technology, Dr. Aiguader 88, Barcelona 08003, Spain; Universitat Pompeu Fabra (UPF). 08003 Barcelona, Spain; Clark University. 01610 Worcester, Massachusetts, United States of America; Barcelona Supercomputing Centre (BSC-CNS). Jordi Girona, 29. 08034. Barcelona, Spain; Institute for Research in Biomedicine (IRB Barcelona), The Barcelona Institute of Science and Technology, Baldiri Reixac, 10, 08028 Barcelona, Spain; ICREA, Pg. Lluís Companys 23, 08010 Barcelona, Spain

**Keywords:** Genome assembly, Heterozygosity, Hybridization, Polyploidy, Aneuploidy

## Abstract

Recent technological developments have made genome sequencing and assembly accessible to many groups. However, the presence in sequenced organisms of certain genomic features such as high heterozygosity, polyploidy, aneuploidy, or heterokaryosis can challenge current standard assembly procedures and result in highly fragmented assemblies. Hence, we hypothesized that genome databases must contain a non-negligible fraction of low-quality assemblies that result from such type of intrinsic genomic factors. Here we present Karyon, a Python-based toolkit that uses raw sequencing data and *de novo* genome assembly to assess several parameters and generate informative plots to assist in the identification of non-chanonical genomic traits. Karyon includes automated *de novo* genome assembly and variant calling pipelines. We tested Karyon by diagnosing 35 highly fragmented publicly available assemblies from 19 different Mucorales (Fungi) species. Our results show that 6 (17%) of the assemblies presented signs of unusual genomic configurations, suggesting that these are common, at least within the Fungi.

## Introduction

Recent developments in high-throughput sequencing and bioinformatic tools have made the process of sequencing the genome of a new organism a routine task for many laboratories. Genome assemblies provide an invaluable resource to understand the biology of an organism at different levels, from the molecular pathways that govern relevant phenotypes to the population structure of its species. The success of a genome assembly is limited by technical aspects as well as by intrinsic properties of the sequenced genome (Gabaldón and Alioto 2016). A successful assembly depends on the quality, design, and depth of the sequencing libraries which must typically adapt to budget limitations. Naturally, if the sequencing methodology or the computational approaches are inappropriate, the results will be poor. However, additional difficulties might arise independently of the methodology employed, due to intrinsic properties of the genome that interfere with genome assembly algorithms.

### Biological factors affecting genome assemby quality

Whole-genome sequencing can be performed using short- or long-read sequencing technologies. The relatively higher throughput, lower error rate and cost still makes short read sequencing the most commonly used approach. Despite the increasing use of long-read technologies for assembly purposes, a large amount of genome assemblies available in public databases have been generated from short reads. Short-read assembly is usually performed using De Bruijn-based assemblers, which are mainly influenced by the number of different *k*-mers (all possible sequences of length *k)* present in the libraries. For this reason, the main intrinsic factors that compromise the success of a genome assembly are the genome size, the sequence heterozygosity, the abundance of low complexity regions (i.e., highly repetitive sequences), as well as the presence of high or uneven ploidy, contaminating sequences or extreme nucleotide compositions (Figure 1).

**Figure 1:**
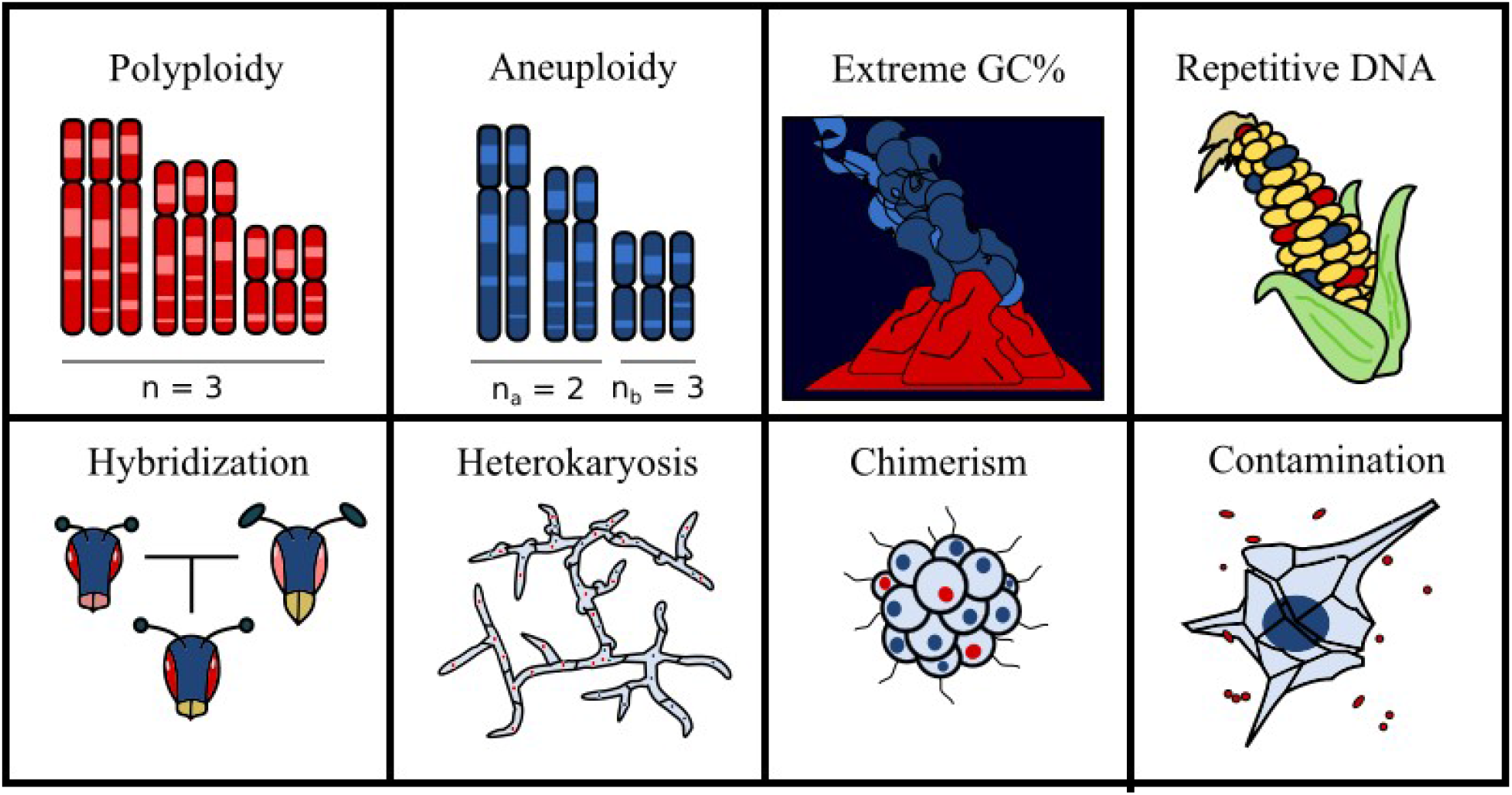
Factors that difficult genome assembly. Ploidy and aneuploidy increase the number of possible states per site. Extreme GC% composition affects the information that different *k*-mers have, and extreme deviations are relatively common in extremophilic organisms. Transposable elements and other forms of repetitive elements increase genome size, affect GC% locally and reduce sequence complexity. Hybridization, heterokaryosis and chimerism introduce two genotypic signals that might be quite divergent, which increases heterozygosity. Finally, contamination introduces undesired sequences with uneven composition, heterozygosity and stoichiometry.

Genome size impacts computational costs, as many assembly algorithms scale non-linearly (Wajid and Serpedin 2012; Simpson and Pop 2015; Wajid et al. 2016). Heterozygosity implies the existence of allelic differences within an individual, which can be single nucleotide polymorphisms, insertion-deletion differences, copy number variations or larger genomic rearrangements. Standard assembly algorithms have difficulty to differentiate between highly heterozygous regions and distinct but highly similar genomic regions (Hirsch and Robin Buell 2013; Leszek P Pryszcz and Gabaldón 2016). This in turn results in fragmented assemblies with inflated size compared to empirical measurements, as many of these regions appear duplicated (Leszek P Pryszcz and Gabaldón 2016), often in short scaffolds. Similarly, repetitive or low complexity genomic regions are difficult to resolve without the aid of expensive experimental approaches, particularly when they span large genomic regions. Duplicated regions introduce multiple possible solutions to the process of scaffolding, increasing assembly fragmentation and computational costs (Hirsch and Robin Buell 2013; Wajid et al. 2016).

Similarly, ploidy deviations can greatly affect genome assembly. The first possible ploidy deviation is polyploidy, which is the presence of more than two chromosomes for the majority of the genome. Polyploidy is generally associated to genome heterozygosity, as it increases the amount of possible states per site (Aguiar and Istrail 2013; Bonizzoni et al. 2016). For a diploid site only two states are possible: heterozygous or homozygous, depending on whether the two alleles are different or equal, respectively. One can differentiate between the two possibilities by detecting frequencies of alternative states higher than expected from sequencing errors, which can be determined with statistical methods. For a triploid, however, there are two possible heterozygotic states, and differentiating between them depends on relative frequencies, which in turn might be affected by stochastic variations in coverage. For a diploid site, the null hypothesis is that the frequency of the alternative site is different from 0. A putative triploid site faces the same hypothesis, but it also needs to test whether it fits better a frequency of 0.5 than a frequency of 0.33 or 0.66 (Weiß et al. 2017). Increasing sequencing depth reduces the effect of stochastic noise, but also increases experimental and computational costs, and the problem is even greater for higher ploidy levels.

Another deviation from the traditional eukaryotic karyotypic organization is aneuploidy, the presence of uneven numbers of chromosomes or large chromosomic regions (Torres, Williams, and Amon 2008; Gerstein and Berman 2015) that do not affect most of the genome. Aneuploidy tends to cause the same problems as polyploidy in assemblies, albeit with the effect being limited only to the aneuploid regions. Because of this, aneuploid regions will tend to appear highly fragmented and might remain undetected if the project is willing to accept a certain level of fragmentation. Genes present in these chromosomes will have a higher likelihood of being unannotated. Animal and plant genomics have traditionally considered aneuploidies as rare events, due to their deleterious effects on many of these organisms, specially during embryonic development. This paradigm is clearly false for many fungal (C. A. Anderson et al. 2015; Berman, Wertheimer, and Stone 2016; Mehrabi, Mirzadi Gohari, and Kema 2017) and protist (Mannaert et al. 2012; Tůmová et al. 2016) lineages, but the lack of traditional cytogenetic studies for many of these organisms makes difficult to have a clear global picture. Eukaryotic genomics has only recently started to focus on pangenomes (Golicz, Batley, and Edwards 2016; McCarthy and Fitzpatrick 2019; Sibbald et al. 2020; Naranjo-Ortiz and Gabaldón 2020; Gerdol et al. 2020), but aneuploidies might be an important confounding factor for these studies.

In syncitial organisms, such as filamentous fungi or slime moulds, there is the possibility of coexistence of genetically different populations of nulei within a cytoplasm, a condition known as heterokaryosis (Maheshwari 2005; James et al. 2008; Strom and Bushley 2016). Heterokaryosis is functionally similar to ploidy, although with some important differences. First, the relative proportions between heterozygous sites do not necessarily adjust to a simple fraction.

In other words, if two populations of nuclei coexist in a cytoplasm, the relative proportion of each type of nucleus is not necessarily equal. Second, since nuclei divide independently from each other, mitotic or meiotic recombination should be rare. This independency implies that any relative chromosomal rearrangements (i.e., duplications, deletions, translocations and inversions) between the two nuclear populations, either pre or post union, would remain in nuclear populations for long periods of time. These rearrangements introduce the aforementioned complications in genome assemblies, and some of these might be difficult to differentiate from other chromosomal aberrations. A similar phenomenon is chimerism, in which the body of an organism is composed by two or more populations of genetically distinct cells. However, in most cases chimeras arise from fusions of two or more embryos and as such the expected effect on heterozygosity is low. Certain lineages, specially colonial species, might arise by fusion of several genetically distinct individuals (Blanquer and Uriz 2011), but very little is known regarding the effect of chimerism in genome assemblies.

The presence of sequence contamination can greatly compromise the quality of the genome assembly (Schmieder and Edwards 2011; Kumar et al. 2013; Trivedi et al. 2014; Laetsch and Blaxter 2017; Lu and Salzberg 2018). Extraneous sequences introduce noise, create chimeric contigs and might introduce errors in *k*-mer estimations. Highly diverse contaminations (e.g. From the gut microbiota) introduce sequences with highly variable level of coverage, heterozygosity and composition; while highly abundant contaminants (e.g. Symbiotic bacteria) are typically more homogeneous in all these parameters, but might still form chimeric contigs and would indirectly reduce the depth of coverage in the main genome. Contaminations reducing the signal of the main genome are particularly problematic for single cell sequencing projects (Huang et al. 2015; Gawad, Koh, and Quake 2016). This is normally prevented by methodological means, but contaminating sequences are intrinsic for certain samples or even organisms, such as the case of symbiotic organisms (e.g. Lichens).

Finally, genomes with extreme compositions, typically very high or low GC content (GC%), can be difficult to assemble. For these genomes, the information contained by AT positions is different than the information contained by GC, as *k-*mers composed of the favoured nucleotide pair will appear at higher frequencies. GC% has a well-documented effect on some sequencing technologies, most notably on the quality of Illumina reads (Benjamini and Speed 2012; Ross et al. 2013). Fortunately, GC% is easy to measure from raw reads, and some genome assemblers include options specially adapted for these cases (Bankevich et al. 2012; D. Scott and Ely 2014). Low GC% is typically associated to high abundance of low complexity regions and transposable elements, but extreme GC% is also a hallmark of certain lineages, such as several groups of early diverging Fungi (e.g. Neocallimastigales, Mucoromycota, Zoopagomycota) (Naranjo-Ortiz and Gabaldón 2019). Despite their effects in genome analyses, GC% in eukaryotic genomes is often ignored. For example, neither NCBI nor Mycocosm report GC% in their assembly information statistics, unlike the Genome OnLine Database (GOLD), which mostly deals with prokaryotic sequences.

If the presence of the factors outlined above is anticipated, specific technical approaches-both experimental and computational-can be used. Contaminating DNA can be identified easily because sequencing coverage, nucleotide composition and phylogenetic signal is usually different from the main genome and several programs have been developed to identify contaminations (Schmieder and Edwards 2011; Kumar et al. 2013; Trivedi et al. 2014; Laetsch and Blaxter 2017; Lu and Salzberg 2018). Ploidy can be easily estimated with cytogenetic techniques, which has been used for animals and plants since the XIXth century. Cytogenetic techniques are time consuming and difficult to interpret for some groups, such as the fungi, due to their smaller chromosomes. Computational approaches exist to estimate composition and ploidy from sequencing reads (Margarido et al. 2015; Mapleson, Accinelli, Kettleborough, Wright, Clavijo, et al. 2016; Weib et al. 2018). Similarly, hybridization can often be detected based on phenotypic traits (intermediate phenotypes and hybrid vigor), an approach that requires phenotypes to observe to begin with. Genomes of hybrid organisms are heterozygous, and some genome assembly software have been designed to be able to handle this situation (Kajitani et al. 2014; Safonova, Bankevich, and Pevzner 2015; Leszek P Pryszcz and Gabaldón 2016), but proper identification of hybrid lineages cannot be done without adequate population and phylogenetic analyses that require additional datasets that might not exist.

Thus, biological factors affecting genome assembly quality increase the overall costs of a project and require expertise that might not be available. Furthermore, such complications are often unforeseen and, if the possible causes for a low-quality assembly are not investigated, it could potentially lead to the direct deposition in databases of low-quality assemblies or even failure of the whole project. Given the difficulty of performing analyses on low quality assemblies, it is likely that published genomes are biased in favor of organisms with canonical genome architectures. As a result, it is so far unknown how common non-standard genome architectures really are.

## Results

### The Karyon toolkit

To aid in the identification of these non-canonical genomic architectures, we developed Karyon, a python-based toolkit that assesses several parameters of sequencing data and their derived assemblies that are common indicators of different intrinsic genomic features that may lead to poor assemblies using standard procedures. Karyon is comprised of different modules that can be used independently or sequentially. Karyon is written in Python 3 and freely available to download as a Docker build or as a standalone project in https://github.com/Gabaldonlab/karyon.

Karyon integrates Trimmomatic (Bolger, Lohse, and Usadel 2014) as an optional step to eliminate low quality positions and adapters from sequencing reads. It then uses that input to generate a *de novo* assembly using SPAdes v3.9.0 (Bankevich et al. 2012), dipSPAdes v3.9.0 (Safonova, Bankevich, and Pevzner 2015), Platanus v1.2.4 (Kajitani et al. 2014) or SOAPdenovo2 v2.04-r240 (Luo et al. 2012). As dipSPAdes was specially designed to deal with highly polymorphic genomes (Safonova, Bankevich, and Pevzner 2015), it was chosen to be Karyon’s default option. Karyon then uses the *de novo* assembly to generate a reduced assembly using Redundans (Leszek P Pryszcz and Gabaldón 2016). Redundans is a pipeline that collapses assembly fragments with high similarity in order to create an artificial haploid genome assembly. This assembly is then used as reference to map the original sequencing reads using BWA-MEM (Li 2013), and generate a variant calling file with GATK v4.1.9.0 (McKenna et al. 2010). A battery of analyses is then performed on the sequencing libraries, the assemblies, and the maps of coverage and genetic variation to generate plots that will aid in the diagnosis of the genomic structure. Figure 2 summarizes the pipeline.

**Figure 2:**
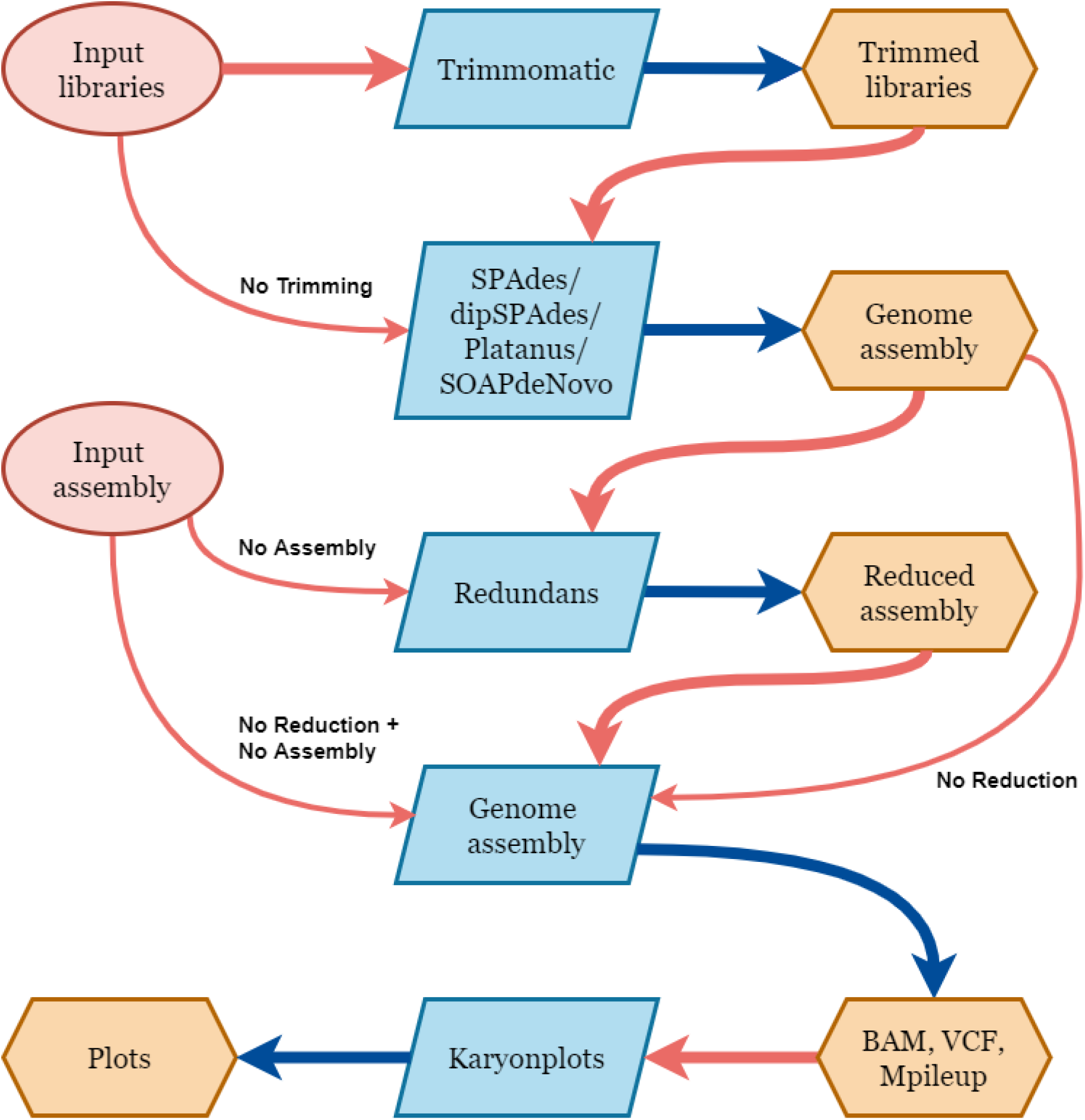
Karyon pipeline. Schematic representation of the steps and program used by Karyon. Red circles represent possible user inputs. Blue boxes represent software used for each step. Orange hexagons represent files generated by the software. Red arrows indicate input to a program, blue arrows represent output of a program. Thicker red arrows represent the standard pipeline, while thinner red arrows represent the different options the user can select to skip some of the steps. These options appear next to the arrow.

Karyon uses the K-mer analysis toolkit (KAT) (Mapleson, et. al. 2016) to provide a *k*-mer (all possible sequences of length *k*) spectrum analysis as part of its report. From this analysis it produces frequency histograms representing coverage versus *k*-mer counts. These plots inform on ploidy and heterozygosity of a genome. In a haploid genome, for *k*-mers of enough size, most *k*-mers will appear either one or zero times, with unique *k*-mers having an average coverage roughly equal to the average depth of coverage. Deviations from these patterns suggest alternative architectures. For instance, the presence of two peaks in the *k*-mer plot may indicate a non-homozygous diploid. To complement these analyses and provide further information on the features of the genome, Karyon assesses scaffold length distributions, relationships between scaffold length and coverage, sliding-window analysis of coverage and genetic variation, as well as allele-frequency distributions per scaffold (Figure 2). In addition, Karyon uses nQuire (Weib et al. 2018) to estimate the likelihood of different ploidy levels in sliding windows per scaffold. Altogether, the interpretation of these analyses can be used to detect polyploidies, aneuploidies, hybridizations, heterokaryosis, large segmental duplications, or the presence of symbiont or contaminating sequences. Further details on each of these analyses and how they can be interpreted are provided in Karyon’s manual. Illustrative, practical examples of their use can be seen in the next section.

Each of the steps is optional and can be controlled with flags in the main script. Additionally, the script uses a configuration file, that allows to define the options of each of the dependency programs. This configuration file is automatically created during the installation and can be modified with any text editor. We encourage the user to make a copy of the original configuration file for future modification. Installation is fully automated, requiring no user input during the process.

### Genomic survey in the Mucorales (Fungi)

To showcase the use of Karyon, we undertook an analysis of deposited fungal genomes in the order Mucorales. Fungi are in a particularly privileged position to assess the impact of non-canonical genomic architectures in genome assemblies. Fungi generally have small and compact genomes and can be often cultured under axenic conditions. As a result, the amount of sequenced fungal genomes is now in the order of thousands, including several strains for many species. Even more, comprehensive efforts to obtain a balanced coverage of the existing fungal diversity are ongoing, such as the 1000 fungal genomes (Grigoriev et al. 2014) and the 1000 yeast genomes initiatives (Wilkening et al. 2013; Strope et al. 2015; Zhu, Sherlock, and Petrov 2016; Peter et al. 2018). Thus, fungi provide an excellent system to study the incidence of different genomic accidents in evolution (Gerstein and Berman 2015; Berman, Wertheimer, and Stone 2016; Todd, Forche, and Selmecki 2017). Despite this, the quality of fungal genomes is often sub-optimal, and databases are riffed with highly fragmented assemblies. Genomic factors such as those discussed above might complicate genome assembly and be responsible for this observed fragmentation, at least partially. Considering this, we hypothesized that genome databases must contain a fraction of low-quality assemblies from fungal organisms that are caused by intrinsic genomic factors. If that is true, reanalysis of the raw data should lead us to describe novel genomic accidents and obtain a minimum estimate of their relative abundance.

We thus applied Karyon to a set of 35 publicly deposited genomes from the fungal order Mucorales. Our results suggest that non-standard genomic organizations are not rare, and that future studies on other groups are likely to uncover many new cases. We selected the order Mucorales because this group comprises several described examples of whole-genome duplication, both at ancient and recent (Ma et al. 2009; Corrochano et al. 2016). Many sequenced members of the clade come from clinical samples, an environment that is known to promote the emergence of different genomic accidents (Schoenfelder and Fox 2015; Todd, Forche, and Selmecki 2017; Mixão and Gabaldón 2018). Additionally, several represented species included two or more sequenced isolates, allowing to get a glimpse at their intra-specific diversity. We obtained 35 genome assemblies from 19 different Mucorales species deposited in GenBank between January 1st 2005 and December 31th 2015 (Table 1). For 4 of the species, dipSPADes was unable to generate an assembly.

**Table 1:** NCBI Assembly statistics for the analyzed strains. Strains with darker background possessed some property that was affecting assembly quality and was diagnosed using Karyon. Fragmentation in all remaining strains is attributed to low sequencing depth.

| Species | NCBI genome size (Mbp) | NCBI number of scaffolds | GeneBank Accession | Genome size after Karyon (Mbp) | Number of scaffolds after Karyon | Diagnosis |
| --- | --- | --- | --- | --- | --- | --- |
| *Rhizopus microsporus* ATCC 62417 | 49.6 | 1386 | GCA_900000135.1 | 40.1 | 5521 | Contamination |
| *Rhizopus microsporus* CBS_344.29 | 49.2 | 1554 | GCA_000825725.1 | 32.1 | 3037 | Contamination |
| *Rhizopus microsporus* B9738 | 75.1 | 5266 | GCA_000697275.1 | 71.6 | 12789 | Misidentification |
| *Rhizopus microsporus* var. *rhizopodiformus* B7455 | 48.7 | 4658 | GCA_000738565.1 | 21.8 | 2176 | Contamination |
| *Rhizopus delemar* Type I NRRL 21789 | 42.0 | 3921 | GCA_000697155.1 | 33.4 | 4824 | Unknown |
| *Rhizopus delemar* Type II NRRL 21446 | 38.9 | 1156 | GCA_000738605.1 | 33.7 | 5071 | Unknown |
| *Rhizopus delemar* Type II NRRL 21447 | 38.7 | 1177 | GCA_000738595.1 | 28.9 | 6683 | Unknown |
| *Rhizopus delemar* Type II NRRL 21477 | 40.8 | 1808 | GCA_000738585.1 | None | None | Unknown |
| *Rhizopus oryzae* 99-892 | 39.1 | 1168 | GCA_000697725.1 | 29.6 | 1875 | Unknown |
| *Rhizopus oryzae* HUMC02 | 40.3 | 2313 | GCA_000697605.1 | None | None | Unknown |
| *Rhizopus oryzae* B7407 | 43.3 | 4683 | GCA_000696915.1 | 34.7 | 3720 | Unknown |
| *Rhizopus oryzae* type I NRRL 13440 | 43.4 | 5022 | GCA_000697075.1 | None | None | Unknown |
| *Rhizopus oryzae* type I NRRL 18148 | 47.5 | 14653 | GCA_000697095.1 | None | None | Unknown |
| *Rhizopus oryzae* type I NRRL 21396 | 42.8 | 4445 | GCA_000697115.1 | 34.2 | 4115 | Unknown |
| *Rhizopus oryzae* 99-133 | 41.5 | 4317 | GCA_000697135.1 | 27.2 | 1332 | Unknown |
| *Rhizopus oryzae* 97-1192 | 42.9 | 4566 | GCA_000697195.1 |  |  | Unknown |
| *Rhizopus stolonifer* B9770 | 38 | 5567 | GCA_000697035.1 | 30.1 | 6406 | Unknown |
| *Mucor circinelloides* B8987 | 36.7 | 2210 | GCA_000696935.1 | 29.9 | 4864 | Unknown |
| *Mucor indicus* B7402 | 39.8 | 3117 | GCA_000697295.1 | 32.1 | 691 | Unknown |
| *Mucor racemosus* B9645 | 65.5 | 6360 | GCA_000697255.1 | 46.0 | 4444 | Hemidiploid |
| *Mucor velutinosus* B5328 | 35.9 | 2411 | GCA_000696895.1 | 28.2 | 2743 | Unknown |
| *Lichtheimia corymbifera* 008-049 | 36.6 | 1629 | GCA_000697175.1 | 42.8 | 3575 | Unknown |
| *Lichtheimia corymbifera* B2541 | 36.6 | 1176 | GCA_000697475.1 | 13.2 | 3575 | Unknown |
| *Lichtheimia ramosa* B5399 | 45.6 | 3968 | GCA_000738555.1 | 26.6 | 861 | Aneuploid, hybrid |
| *Saksenaea oblongisporus* B3353 | 40.8 | 1702 | GCA_000697495.1 | 29.7 | 622 | Unknown |
| *Saksenaea vasiformis* B4078 | 42.5 | 2417 | GCA_000697055.1 | 32.7 | 1506 | Unknown |
| *Cokeromyces recurvatus* B5483 | 29.3 | 2637 | GCA_000697235.1 | 26.6 | 5213 | Unknown |
| *Syncephalastrum monosporum* B8922 | 29.6 | 1284 | GCA_000697355.1 | 24.1 | 5271 | Unknown |
| *Syncephalastrum racemosum* B6101 | 29.6 | 1035 | GCA_000696955.1 | 23.3 | 311 | Unknown |
| *Cunninghamella elegans* B9769 | 31.7 | 1380 | GCA_000697015.1 | 30.8 | 5465 | Unknown |
| *Apophysomyces elegans* B7760 | 38.5 | 1528 | GCA_000696995.1 | 29.3 | 1293 | Unknown |
| *Apophysomyces trapeziformis* B9324 | 35.8 | 1400 | GCA_000696975.1 | 30.1 | 898 | Unknown |
| *Thermomucor indicae-seudaticae* HACC 243 | 29.6 | 1958 | GCA_000787465.1 | 25.7 | 4118 | Unknown |
| *Parasitella parasitica* CBS 412.66 isolate NGI315 ade-mutant | 44.9 | 15637 | GCA_000938895.1 | 23.5 | 3295 | Unknown |

Karyon was run using the complete default pipeline. Most of the analyzed genomes (27, 79.4%) presented very low levels of heterozygosity and a relatively homogeneous coverage across the genome, suggesting that those strains are haploid or, if presenting higher ploidy, extremely homozygous. Fragmentation in these cases might be caused by insufficient coverage, presence of repetitive regions or some other methodological constraints. However, our pipeline uncovered cases that produced anomalous results in the different Karyon tests. Below we describe these cases and propose a plausible scenario to explain each of the obtained results based on the data obtained from the Karyon pipeline.

### *Rhizopus microsporus* species complex

At the moment of this study, eight *Rhizopus microsporus* strains were deposited in the NCBI database. Interestingly, three of them presented a genome size estimated around 25Mbp; four of them had a genome size close to 50Mbp; and one presented a genome size of 75Mbp. Only the three strains with a genome size of 25Mbp had sufficiently good assemblies considering they were based on short read, with a scaffold number below 1000, and thus were not selected for further analyses. Additionally, the raw libraries for one of the strains presenting 50Mbp genome assembly size *(Rhizopus microsporus* var. *chinensis* CCTCC M201021) were not publicly available and thus could not be part of the survey. For the remaining three strains with genome size close to 50Mbp (ATCC62417, CBS344.29 and var *rhizopodiformis* B7455), our *de novo* assembly pipeline recovered a genome size of approximately 40Mb, which is smaller than the assemblies deposited in NCBI (Table 1). The heterozygosity distribution in these assemblies shows that most of the genome presented a relatively uniform behavior with low heterozygosity. In all three cases, though, a considerable proportion of the genome appears with a highly variable coverage and increased heterozygosity (Figure 3). For these three strains, BlobTools (REF) shows an important fraction of the genome which seems to be of bacterial origin (Figure 3b) and thus we conclude that contamination is the main cause of assembly fragmentation.

**Figure 3.**
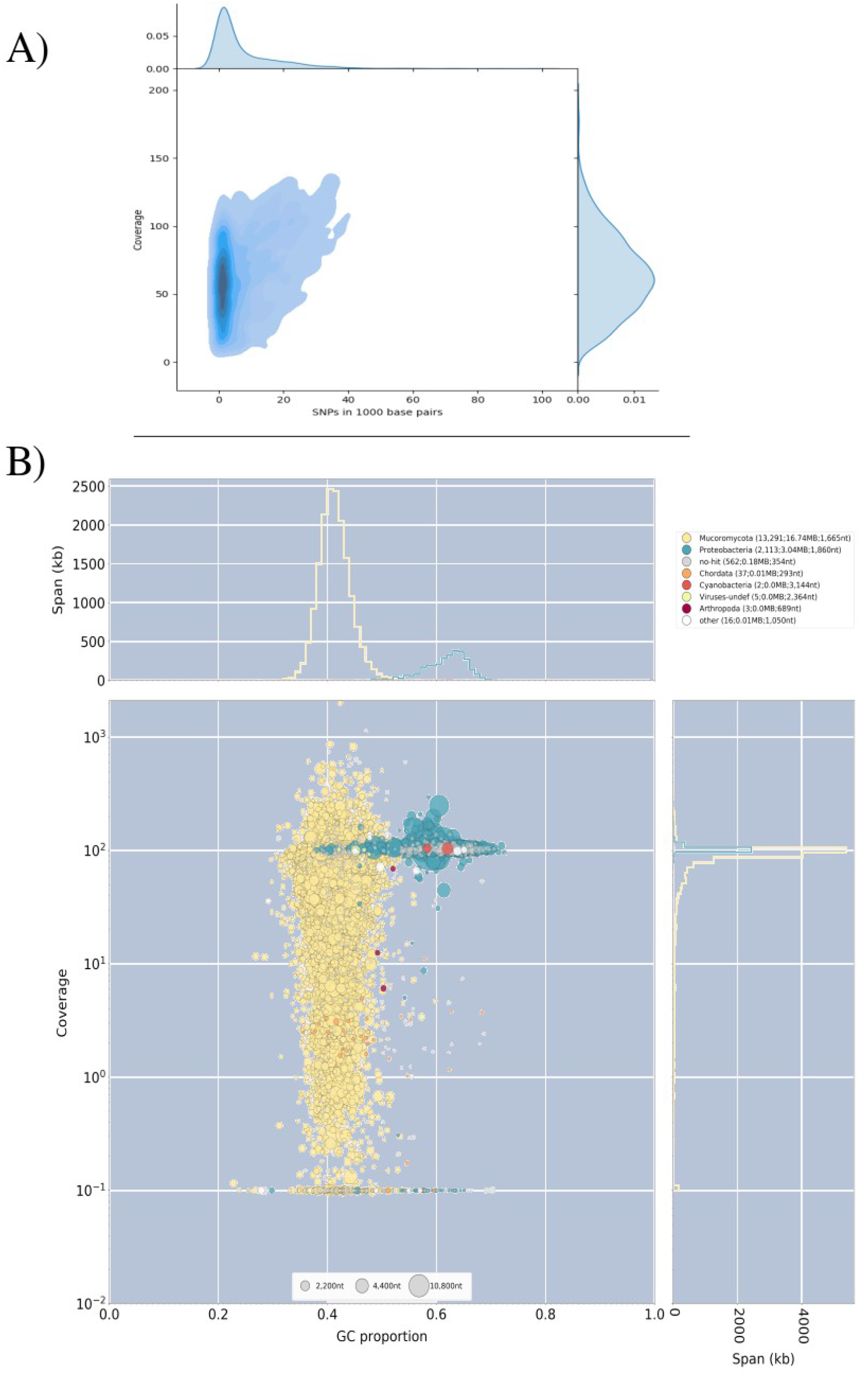
Analysis of *Rhizopus microsporus* ATCC62417. A) Variation versus coverage plot reveals the existence of a highly variable portion of the genome that presents variable heterozygosity levels. B) BlobTools analyses suggest that the genome presents a considerable portion of contaminating sequences. Coverage of the sequences assigned to bacteria is very low when the analyses are performed with other libraries (Data not shown), which proves that the conflicting signal observed in this sample has its origin in a contaminated sequencing library. Results for *R. microsporus* CBS344.5 and var. *rhizopodiformus* B7455 show similar patterns of contamination (data not shown).

The remaining strain, B9738, showed a surprisingly large genome size in both the assembly deposited in NCBI (75Mbp), and the one reconstructed here (71Mbp). The genome of *R. microsporus* B9738 presents an extremely low level of heterozygosity and a very homogeneous coverage. *K*-mer spectrum also shows just one very clear peak. All in all, all this suggests that B9738 is haploid, despite presenting a 3-fold increase in genome size as compared to other strains of the same species (Figure 4). Augustus gene prediction returned a total of 21,300 gene models, which is an unusually large number for a filamentous fungus. As a reference, the seven genomes in the Rhizopodaceae, to which *Rhizopus* belongs, available in Mycocosm range from 25 to 46 Mbp and from 10,781 to 17,676 annotated genes. Contamination analysis does not suggest the presence of widespread contamination that could explain such over-inflated genome (Figure 4). For this reason, we suggest that B9738 might be a misidentified strain that does not belong to the *R. microsporus* species complex. Indeed, phylogenomic analyses recover B9738 as sister to a clade containing *Mucor* and *Parasitella,* rather than allied with the rest of the *Rhizopus microsporus* species clade (Figure 5), thus supporting a misidentification. It is noteworthy that no sequenced species of either *Mucor* or *Parasitella* have genomes above 49Mbp or with more than 15,000 genes, at least from the available genomes in Mycocosm.

**Figure 4.**
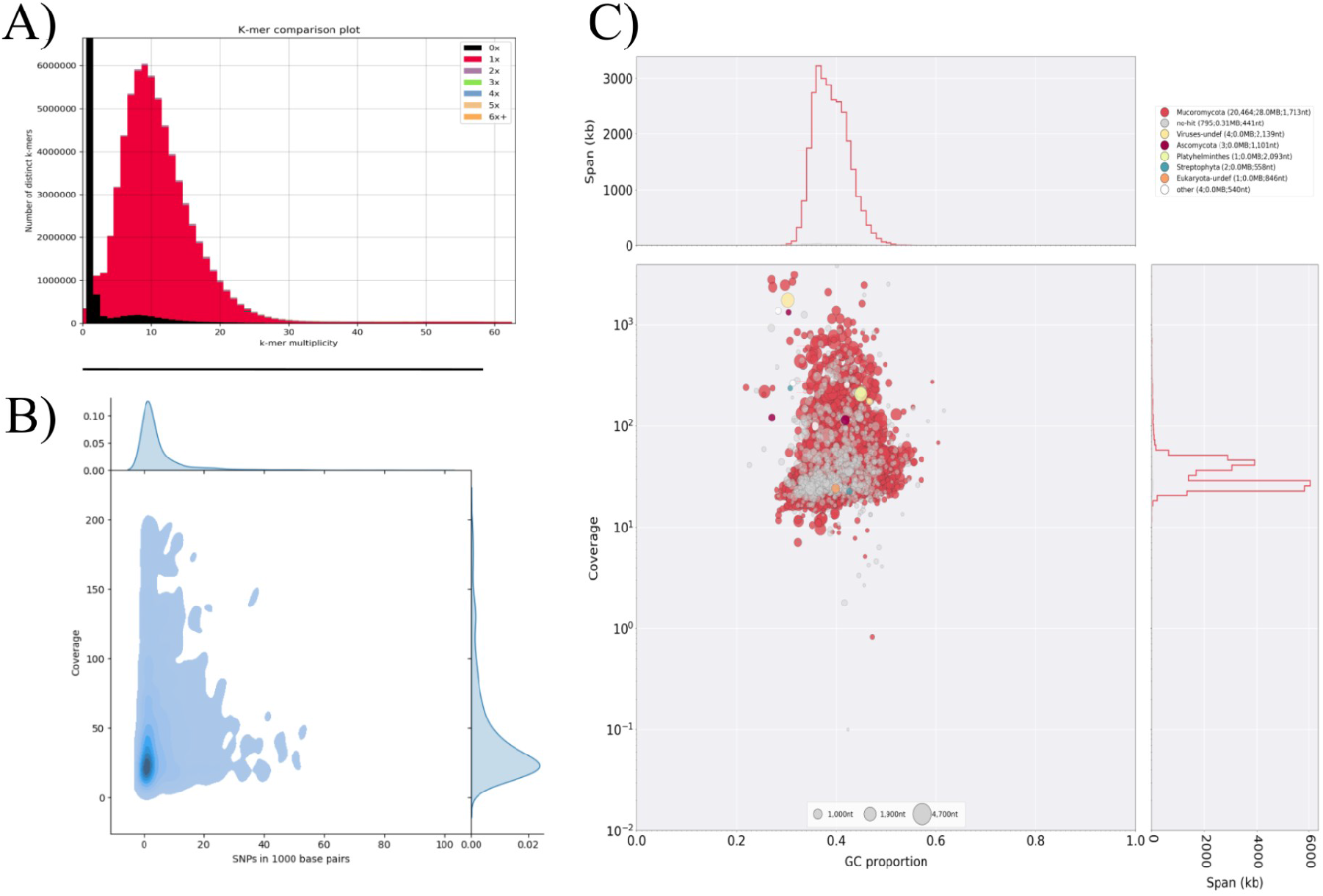
Analysis of *Rhizopus microsporus* B9738. **A)** KAT *k*-mer plot shows very low genome compaction (black area), suggestive of a haploid genome. **B)** Variation versus coverage plot reveals a single main behavior for the genome with regards of its SNP density and coverage. **C)** BlobTools analysis shows no sign of widespread contamination that might be inflating the genome.

**Figure 5.**
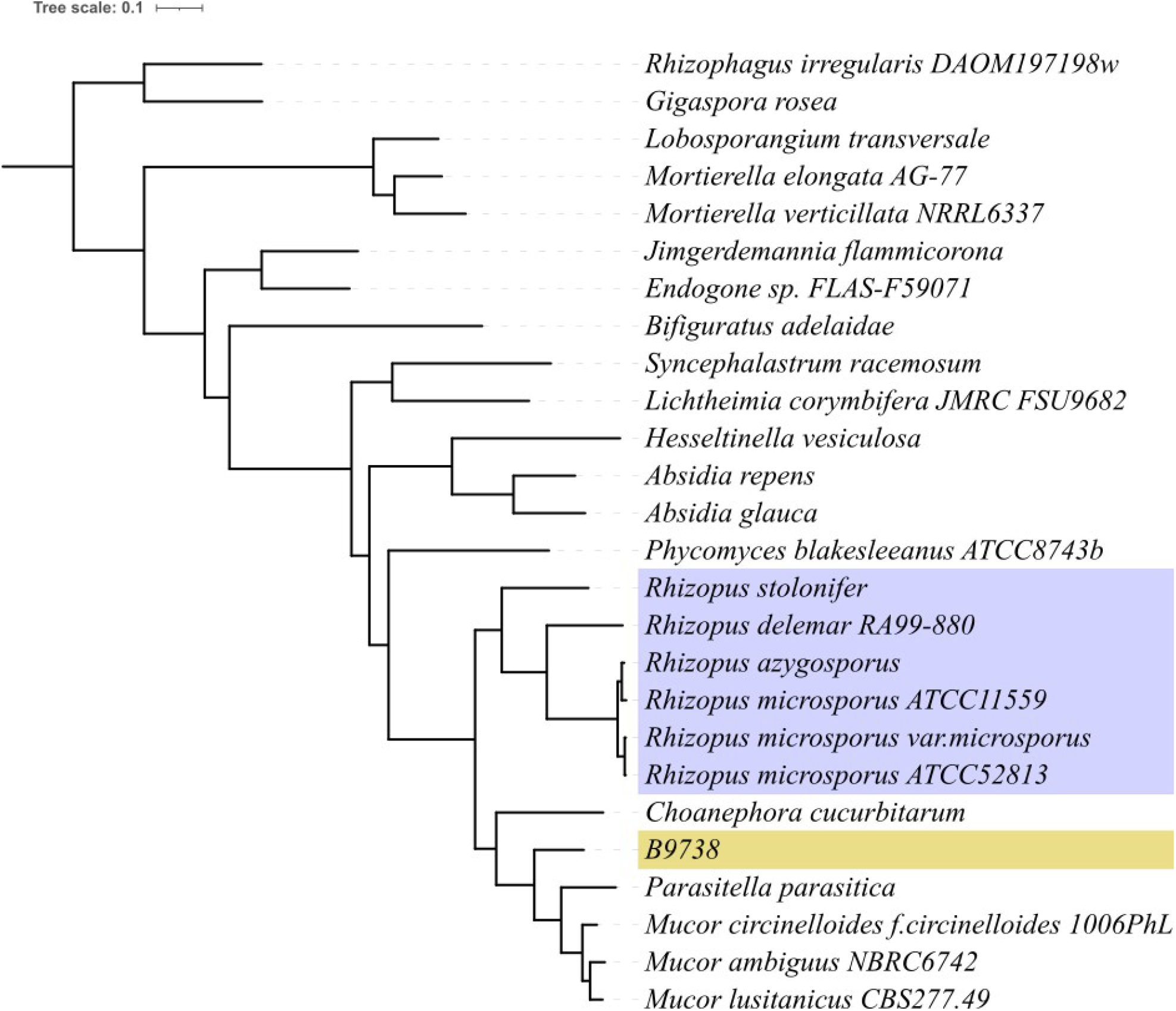
Phylogenetic tree of *Rhizopus microsporus* B9738. Phylogenetic tree inferred from OrthoFinder. The *Rhizopus microsporus* species complex is marked in blue. The problematic strain, B9738, is marked in yellow.

### *Mucor racemosis* B9645

Analyses on *Mucor racemosus* B9645 depicted a genome with a dual behavior. The distribution of heterozygosity and coverage showed two peaks with very low heterozygosity but with different coverage (Figure 6b). This was further confirmed by the *k*-mer spectrum analysis, which revealed two clear peaks (Figure 6a). The genome available in NCBI is 65.5Mbp-long, noticeably larger than the 45.9Mbp we recovered in our analyses (Table 1). The reduction step of Redundans cannot explain this difference, as the assembly size prior to this step is already 46.8Mbp, very close to the final result. Our analyses suggest that contaminating sequences are very minor and do not explain the observed pattern (Figure 6). We hypothesize that *M. racemosus* B9645 is a hemidiploid, which presents a portion of its genome in haploid state, and other portion in a highly homozygous diploid state. Due to the low heterozygosity exhibited by this strain, the observed genome architecture might have arisen by either autopolyploidization followed by chromosome loss or by chromosomal duplications.

**Figure 6.**
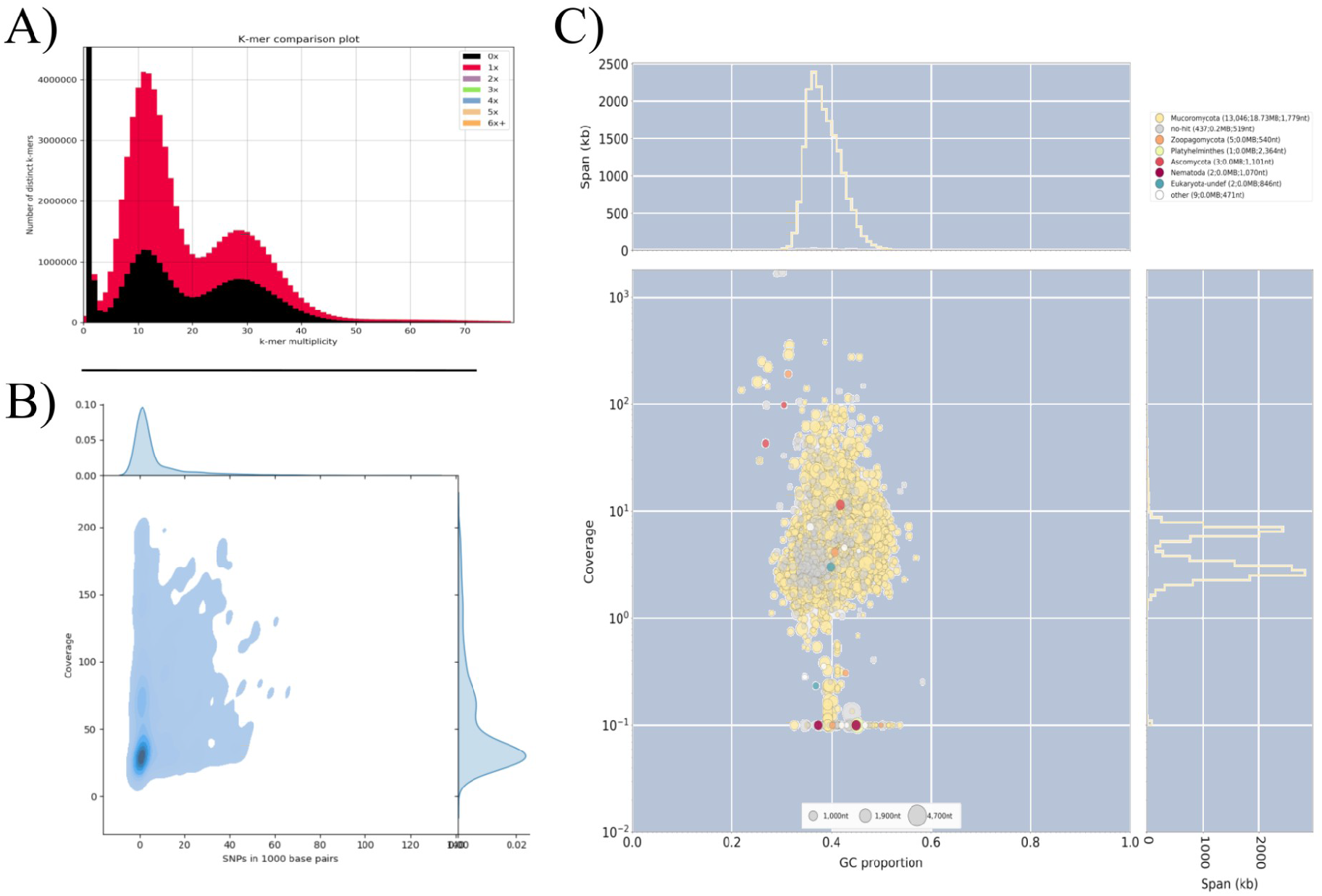
Analysis of *Mucor racemosus* B9645. **A)** KAT *k*-mer plot shows two peaks of coverage considerably affected by genome reduction (black area), suggestive of a highly heterozygous diploid genome. **B)** Variation versus coverage plot reveals a bimodal behaviour for the genome with regards of its coverage, but both peaks appear with very low SNP density. **C)** BlobTools analysis shows no sign of widespread contamination that might be inflating the genome.

### *Lichtheimia ramosa* B5399

The Karyon assembly for this genome was only 26.6Mbp, much smaller than the NCBI assembly (45.6Mpb long, Table 1). Unlike other genomes, our assembly presented a considerable improved quality, going from 3,968 scaffolds and N50 of 33,650 in the NCBI assembly to 861 scaffolds and N50 of 133,635 in our own assembly. *L. ramosa* presents a heterozygosity level around 3% in its diploid peak (Figure 7). All considered, we propose that *L. ramosa* B5399 is a diploid with high heterozygosity and several aneuploid (both aneuploid and triploid), likely resulting from mating between two distantly related strains, and the NCBI assembly is inflated as a consequence of this situation.

**Figure 7.**
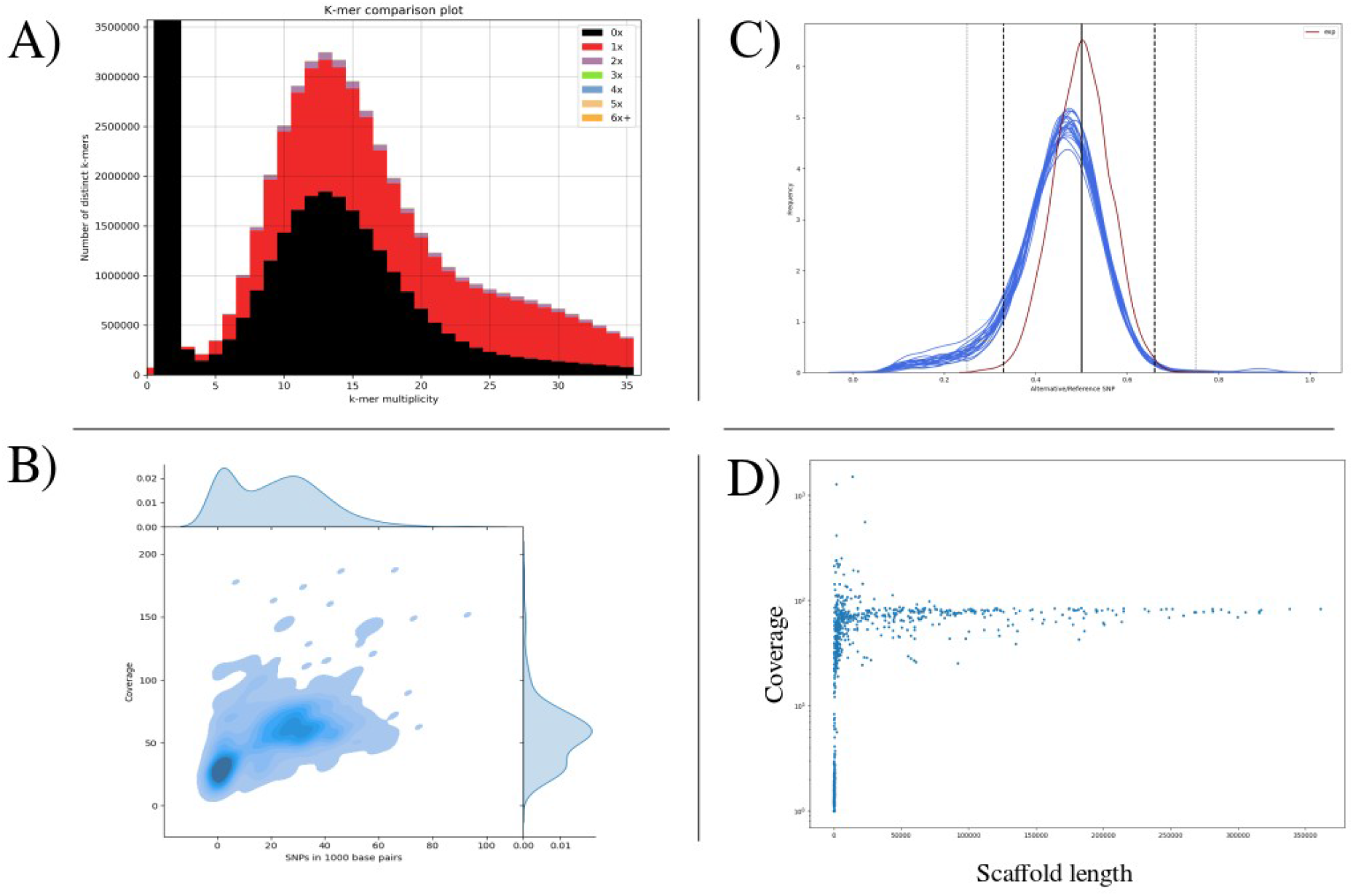
Analysis of *Lichtheimia ramosa* B5399. **A)** KAT *k*-mer plot shows one peak with considerable genome compaction (black area) suggestive of a diploid genome. **B)** Variation versus coverage plot reveals a unimodal behaviour for the genome with regards of its coverage, presenting a widespread heterozygosity of approximately 3% (maximum density around 30 SNP/Kbp). **C)** Alternative allele frequency shows that all scaffolds present a behaviour very similar to the ideal diploid. **D)** Scaffold length plot shows that, with the exception of a group of very low coverage scaffolds, all the genome presents a uniform coverage.

## Methods

### Sequencing data

We downloaded raw data from libraries deposited at Short Read Archive (SRA)(National Centre for Biotechnology Information 2015) of those species in the Mucorales with a highly fragmented assembly (>1,000 scaffolds), which included at least one paired-end Illumina library larger than 1Gb after quality filtering (Table 1), to ensure at least a decent coverage. Since most of our genomes have typical assembly sizes around 40Mbp, this measure ensures a bare minimum average coverage of 20.

### *De novo* gene annotation

We used Augustus v3.1.0. (Brudno et al. 2003) to obtain a *de novo* gene prediction using the included *Rhizopus oryzae* trained model.

### Contamination detection

For each of the conflictive assemblies, we generated an Augustus prediction. Then, we used Blastp (Stephen F. Altschul, Warren Gish, Webb Miller 1990) to query the whole proteome against Uniref100 (Consortium 2014). Since the genomes come from public databases, their own proteins should appear as hits and thus we retrieved the 10 best hits. We have used these hits to assign a taxonomic profile. Additionally, we have used the predicted Augustus CDS to map sequencing reads with GATK. With both the taxonomic profile and the variant calling file, we have run BlobTools (Laetsch and Blaxter 2017) in order to identify the presence of widespread contamination in the sequencing libraries.

### Phylogenomic analyses

In order to identify the phylogenetic position of *R. microsporus* B9738 we used the Augustus gene prediction and the proteome of 24 other zygomycetes to run OrthoFinder v.2.3.3 (D.M. and S. 2018) with the flags -S blast and -m msa.

## Discussion

As genome sequencing has moved away from model organisms, it has become apparent that many possible genomic architectures are possible, and many do exist in a wide range of organisms. Most of these genomic accidents are difficult to identify from sequencing data alone. As far as we know, Karyon is the first software developed with the intention of analyzing the presence of such genomic incidents during the process of *de novo* genome assembly. We have designed this software to be easy to install and use, with the possibility of installation from both GitHub and Docker.

Despite the success in the implemented strategy, we consider our software has several limitations. Karyon requires an assembly step and variant calling protocol, for which some default options are included. However, the included programs might not suit every need. For example, extremely large genomes might require alternative assemblers that are not included in our pipeline, or some users might prefer a different set of programs for the variant calling protocol. For those cases Karyon can be used as independent steps (Figure 2). At this moment, the pipeline assumes the use of at least one Illumina paired-end sequencing library. Because of this, we recommend the use of other genome assemblers if other sequencing technologies (i.e., Nanopore or PacBio long reads) are to be used, and the same goes for variant calling protocols. Fortunately, thanks to the modular nature of Karyon, implementation of new programs is straightforward.

We provided a practical example of the usage of Karyon on a publicly available set of fungal genomes from the order Mucorales. While the majority of analyzed assemblies show no sign of any of the considered biological conditions, we were able to effectively find underlying nonstandard genomic architectures that had been previously unnoticed in these assemblies. These results suggest that many authors do not take into consideration this kind of genomic accidents, which in turn greatly hampers the results that might be obtained from them.

How common are these non-standard genomic architectures? Our results suggest that they might be quite abundant, although, so far, they are restricted to a limited selection of species within a narrow clade of Fungi. As such, these genomic anomalies might, or might not, be common in other lineages. However, we consider that there are three important arguments in favor for considering our dataset an underestimation of the abundance of unorthodox fungal genomes, even within the taxonomic range we have selected. The first one is the fact that fungal biomass used for DNA extraction and subsequent sequencing typically comes from cultures. This implies an important ecological step in which the fungus grows at optimal speed and in the absence of most stressors. Aneuploidies, polyploidies and other similar genomic rearrangements are common in the presence of stressors (C. A. Anderson et al. 2015; Berman 2016; Berman, Wertheimer, and Stone 2016; Todd, Forche, and Selmecki 2017), but seem to be out-competed by euploid cells under optimal conditions (Kumaran, Yang, and Leu 2013; Zörgö et al. 2013; A. L. Scott et al. 2017). Hence, isolates growing in rich medium will be selected to lose most chromosomal aberrations they might present. Analogously, many of these chromosomal aberrations might exist in nature but are unable to grow on optimal medium. The advance of environmental sequencing and single cell based technologies might cast some light in this matter in coming years. Supporting this argument, Ahrendt et al. sequenced several environmental isolates of zoosporic and zygomycetous microfungi using these techniques and found several aneuploids and polyploids (Ahrendt et al. 2018). The frequency of unconventional genomic architectures is very likely lineage-dependent. While some of these are well known, such as the dikaryotic phase in Agaricomycetes or the macro and micronuclei of ciliates, strange genomic architectures might be common in more obscure lineages. This not only represents a yet-to-know facet of the biology of these organisms, but it could potentially complicate their study. The third factor to consider is purely human. The datasets we have analyzed were uploaded by researchers who considered they were good enough to be uploaded to a public repository. Thus, it is to be expected that many more low-quality assemblies would have never been never deposited and sit forgotten in the disks of laboratory computers, if not discarded completely.

Even if we consider these possible biases as negligible, our results recover a significant fraction of publicly available genomes with unorthodox genomic configurations. These have been correlated in many fungal groups with adaptation to novel environments (Lenassi et al. 2013; Kravets et al. 2014; Sinha et al. 2017), resistance to antifungals (Harrison et al. 2014; M. Z. Anderson et al. 2017), pathogenic capabilities toward both animals (W. Li et al. 2012; Morrow and Fraser 2013; Leszek Piotr Pryszcz 2014; Gerstein et al. 2015; Mixão and Gabaldón 2018) and plants (Garbelotto et al. 2004; Depotter et al. 2016) and adaptation to industrial settings (S. a. James et al. 2005; Louis et al. 2012; Borneman et al. 2014; Walther, Hesselbart, and Wendland 2014; Peter et al. 2018; Avramova et al. 2018). Beyond that, contamination in sequencing libraries is a problem that can affect any assembly project and might mislead downstream inferences if left unaddressed. Validation of published results goes far beyond the interest of discovering overlooked findings. Comparative genomic studies are limited in their scope and reliability by the quality of assembly and annotation of the genomes, factors that can be greatly compromised by these biological factors. Comparative studies commonly require the use of flagship genomes that represent a given taxon. Often, this generates a chronology of comparisons versus the reference that shapes the perspective on the group. As such, artifacts and errors in strategic genome assemblies, such as reference strains or strains in groups with few represented species, might have a domino effect impacting future studies. Long-read sequencing technologies, which are increasingly being used for genome assembly projects, hold the promise of providing much more information that could be used to resolve many of these unorthodox genomic architectures. However, these approaches require novel computational approaches to fully employ their potential.

## Findings

- We present Karyon, a python-based bioinformatic pipeline that integrates genome assembly and a series of structural analyses for the diagnosis of problematic genomic structures. Karyon is freely available in github and as a docker container (https://github.com/Gabaldonlab/karyon).
- We applied Karyon to 35 highly fragmented, publicly available genome assemblies to identify putative undescribed deviations in genomic architecture that might have caused problems in a standard assembly process. From 35 assemblies, six presented features that suggested possible underlying biological factors as the likely cause of the observed assembly fragmentation. Even though our sample size is small and restricted to a single lineage (Mucoromycotina), our results suggest that the number of unreported deviations in genome architecture in Fungi is considerable. This is emphasized if we consider that most researchers that have produced low quality assemblies are unlikely to publish their data.

## Acknowledgements

TG group acknowledges support from the Spanish Ministry of Science and Innovation for grant PGC2018-099921-B-I00, cofounded by European Regional Development Fund (ERDF); from the Catalan Research Agency (AGAUR) SGR423; from the European Union’s Horizon 2020 research and innovation programme (ERC-2016-724173); from the Gordon and Betty Moore Foundation (Grant GBMF9742) and from the Instituto de Salud Carlos III (INB Grant PT17/0009/0023 – ISCIII-SGEFI/ERDF).

